# Nutrient deprived growth of *Streptomyces* promotes foraging growth and enhanced antimicrobial activity

**DOI:** 10.1101/2024.04.15.589617

**Authors:** Niklas Söderholm, Hugh Tanner, Linda Sandblad

## Abstract

Species of the genus *Streptomyces* are filamentous bacteria mainly residing in soils. The capacity of streptomycetes to adapt to various environments is reflected in the sheer size of their genomes and the number of encoded regulatory proteins. The nutrient-rich laboratory conditions used for culturing significantly differ from the conditions in the natural habitat of streptomycetes and fail to stimulate their full range of adaptive behaviors; as on of the results, the majority of biosynthetic gene clusters remain silent. Moreover, the use of rich media has led to the assumption that nutrient depletion ultimately promotes sporulation. However, we demonstrate that *S. coelicolor* colonies can respond to nutrient depletion by displaying a previously unidentified phenotypic transition termed “foraging” due to its submerged and continuous growth on depleted media. The foraging phenotype is distinctly different from conventional phenotypes in terms of colony morphology, genomic stability, and metabolomic profile. This adaptation to nutrient deprivation through foraging is found throughout the *Streptomyces* phylogeny, indicating that the phenotype is highly conserved. Furthermore, foraging *S. coelicolor* gains the ability to inhibit molds and enhanced competitive activity against both gram-negative and gram-positive bacteria is detected in other species. These findings highlight how morphological adaptations of streptomycetes in nutrient-limited environments alter secondary metabolite production, enabling the screening of novel antimicrobial activities. These discoveries have implications ranging from the basic biology of streptomycetes to drug discovery and microbial ecology.

## Introduction

The gram-positive genus *Streptomyces*, known for its filamentous morphology and production of bioactive secondary metabolites, plays a crucial role in soil as saprophytes that hydrolyze a wide array of polysaccharides and macromolecules [1]. Streptomycetes undergo a complex life cycle, beginning with spore germination, progressing through vegetative hyphal growth with branching to form a compact mycelial network, giving rise to the convex colony appearance. A differentiation process leads to the development of aerial hyphae and spores, which often coincides with production of many secondary metabolites. The differentiation relies on the *bld*-genes to control the aerial development phase and the *whi*-genes for spore maturation [2].

One of the *bld* genes, *bldA* encodes a Leu-tRNA^UUA^ necessary for translation of mRNA UUA codons [3–6] and hence regulates the expression of genes containing this rare leucine codon. One of the genes under this regulation is *adpA*, a global regulator of development and secondary metabolites [7–11]. Interestingly, AdpA also controls the transcription of *bldA* [12]. The life cycle of streptomycetes has been extensively studied on rich media, where subsequent decreasing nutrient levels are believed to promote mycelial differentiation and secondary metabolite production. In a study using rich media, it was shown that *S. coelicolor* metabolite production was enhanced because of genomic instability leading to a genomic heterogeneity and a consequent lower fitness reduction and eventual extinction of these cells [13, 14].

While the importance of optimized growth media for metabolite production and life cycle progression is widely recognized [15, 16], little is known about streptomyces growth and metabolite production under nutrient poor conditions. An observation that aligns with its adaptation to a life as a saprophyte in the soil, is the abundance of gene regulatory proteins, two-component systems, transporters, and substrate-binding proteins, as first identified in the *S. coelicolor* genome, suggesting a significant capacity for adaptation and the utilization of varied nutrient sources [17, 18]. Co-incubation of certain streptomycetes with yeast on rich media lead to the discovery of the “exploratory phenotype”, which in turn led to the understanding that specific carbon sources alone could induce this phenotype [19, 20].

The exploratory growth phenotype, like conventional growth, is connected to metabolite production and it was discovered that production of cryptic metabolites was activated [19]. Identification of biosynthetic gene clusters (BGCs), the genes coding for proteins synthesizing the metabolites, has revealed that each *Streptomyces* species could have the capacity to produce over 20 secondary metabolites [17, 21–23]. However, current standard cultivation methods often fail to trigger the regulatory pathways responsible for their synthesis [24, 25] and only a fraction of possible metabolites have consequently been studied for biological activities. Rapidly consumed carbon sources often repress production of secondary metabolites [26]. However, after primary substrates have been exhausted, carbon catabolite repression adapts growth and metabolism to utilize less prioritized substrates. Therefore, altering environmental stimuli to activate metabolite production from cryptic BGCs is a promising strategy in the search for new bioactive compounds [27].

Considering the saprophyte niche of streptomycetes in soil environments, we sought to determine how growth is affected when degradation of polysaccharides becomes a necessity because it is the main source of carbon. This represents an environment that, in terms of carbon sources, would better reflect their *in situ* growth.

We demonstrate that global nutrient depletion in growth media coincides with a morphological transition of *S. coelicolor* colonies to a phenotype termed “foraging”. Observations throughout the phylogeny suggest that foraging is common to all streptomycetes and also present in a closely related genus. We explore this novel growth morphology, investigate antimicrobial properties, and compare metabolomic profiles of conventional and foraging growth. This study indicates that nutrient-deprived growth may be key to both discovering novel bioactive secondary metabolites and providing a more complete understanding of the *Streptomyces* life cycle.

## Results

### Phenotype transition correlate with nutrient depletion

To investigate how *Streptomyces* colonies respond to nutrient depletion, *S. coelicolor* was incubated on different diluted agar media. A unique growth morphology of *S. coelicolor* was observed after 60 days of incubation on 0.5x diluted tryptone soya agar (TSA). The colonies had developed a thin, veil-like, outer growth zone, distinct from the conventional raised *S. coelicolor* morphology (Figure 1A). We further investigated this morphology, termed “foraging”, using time-lapse imaging over 75 days, which revealed the phenotype transition around day 50 (Video S1).

**Figure 1.**
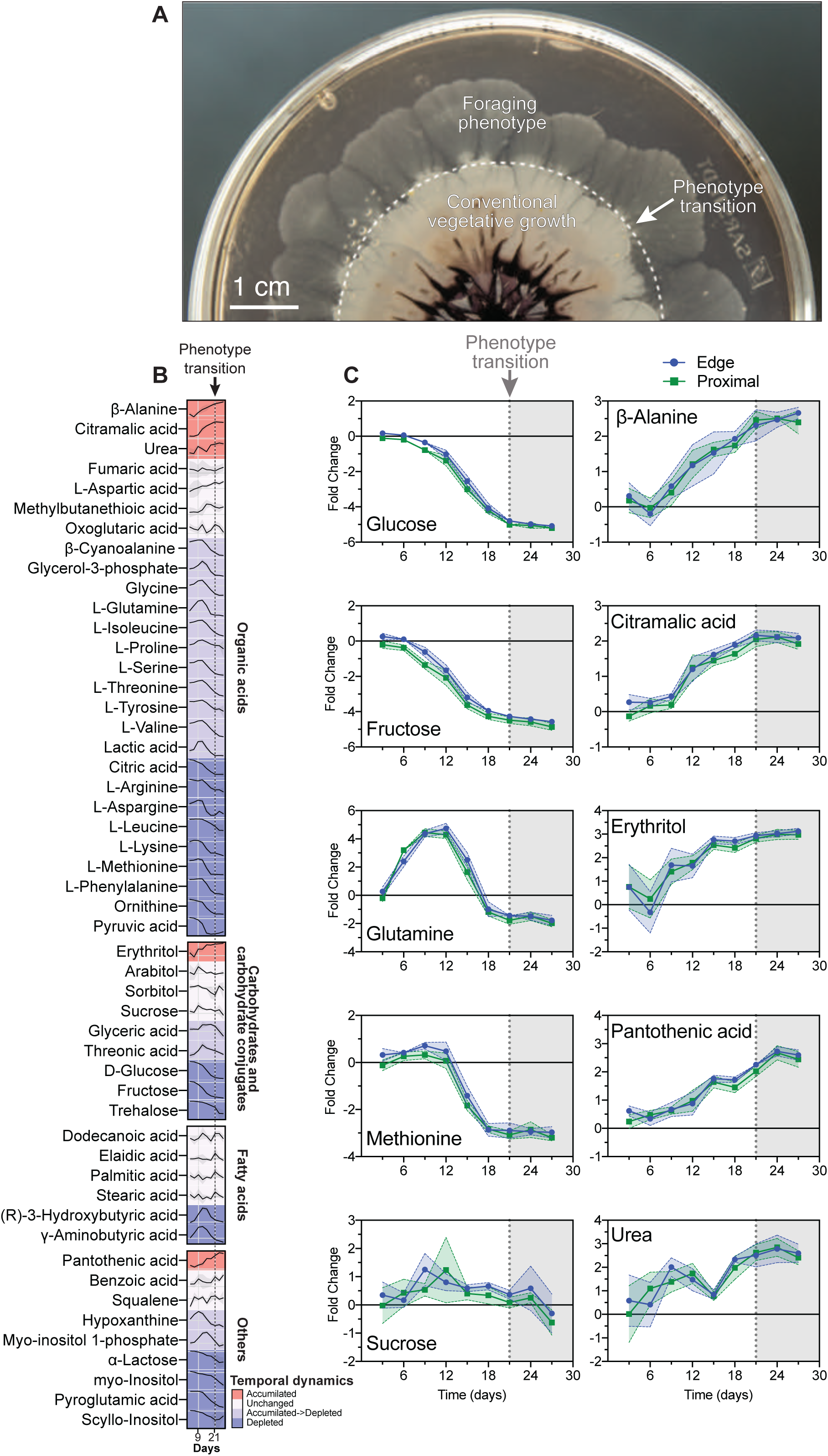
*S. coelicolor* growth phenotype transition correlates with nutrient depletion. (**A**) Colony appearance after 75 days growth on 0.5xTSA. The point of phenotype transition is indicated by the dotted circle. (**B**) Line plot visualization comparing nutrient proximal nutrient levels during *S. coelicolor* growth on 0.5xTSA. Each nutrient is categorized by color to indicate its trend throughout the time series. The measurements proximal to the colony were taken in hexaduplicate for each time point and normalized against the controls. Day 21 when the transition to foraging growth was completed is indicated with the black dotted line. (**C**) Examples of metabolites levels throughout the study. The time point where the transition to foraging growth was observed is indicated with the dotted lines on day 21. Each measurement proximal to the colony and at the edge of the dish hexatuplicate were taken for each time point and normalized against the controls.

It became clear that foraging growth was linked to nutrient availability, as it occurred earlier on diluted media, in smaller media volumes, and was delayed on richer media. The foraging transition was evident already after 21 days when grown in smaller ⌀ 4 cm dishes with 0.5xTSA. Using these smaller dishes, we monitored nutrient depletion by inoculating plates with spores and sampled the agar every three days for 27 days to obtain time point both pre- and post-transition. Agar samples were taken from two locations: (1) 5 mm away from the colony edge to ensure that the sample was not contaminated with hyphae, and (2)from the very edge of the dish, which represents the farthest possible point from the colony. The nutrient levels in the agar samples were analyzed using mass spectrometry. Altogether, 51 metabolites were annotated, including sugars, amino acids, and fatty acids, most of which depleted over time (Figure 1B). Nutrient levels proximal to the colony mirrored those at the dish-edge (Figure S1A). These observations are consistent with calculations based on agar diffusion coefficients that suggest nutrients like e.g., glucose can diffuse 1.5 cm^2^ over three days [28]. This explains the persistence of the foraging phenotype even as it expands into new areas of the agar.

Although the levels of certain metabolites, such as glucose and fructose, were decreasing from the start and depleted at the onset of foraging, others, such as glutamine and lactic acid, initially increased before decreasing. Some metabolites, such as leucine and phenylalanine, remained stable until day 12 before decreasing (Figure 1C, Figure 1, Figure S1B). The levels of certain metabolites, such as sucrose, remained unchanged throughout, and only five metabolites were observed to increase more than one-fold (Figure 1C). Supplementation of 0.5xTSA and MS agar with the metabolites detected to accumulate during foraging-transition did however not induce foraging growth, suggesting they do not act as signals for phenotypic transition.

We conclude that the observed phenotypic transition is nutrient dependent and that the presence of polysaccharides in the form of the agar matrix does not interfere with the foraging phenotype.

### Absence of easily accessible nutrients enables foraging *S. coelicolor* growth

The transition to the foraging phenotype coincided with media depletion; hence the growth of *S. coelicolor* was next investigated in the absence of excessive nutrients. Even the nutrient-depleted media clearly contained complex carbohydrates in the form of agar at the onset of foraging. Hence, an agar medium without additional carbon sources was sought. *Streptomyces* minimal medium contains only three carbon sources: agar, L-asparagine, and glucose [29]. Substituting L-asparagine with ammonium sulfate and excluding glucose from the medium made agar the sole carbon source; this medium is hereafter referred to as foraging agar medium (FAM). Colonies on FAM germinated and displayed similar characteristics to those on 0.5xTSA after the phenotypic transition, exhibiting a thin, “veil-like submerged expansion” that eventually covered the entire plate (Figure 2A). Over time, sectors developed within the colony, some of which became pigmented (Figure 2B). Aerial hyphae formation and pigment production at the inoculation site were influenced by the presence of metabolizable nutrients such as glycerol present in the spore storage buffer (Figure 2C), further demonstrating the effect of nutrients on the foraging phenotype. FAM supplemented with glucose or fructose effectively inhibited foraging growth, leading to conventional raised colonies (Figure S2A). Notably, sucrose supplementation of FAM did not prevent foraging growth, indicating the inability of *S. coelicolor* to metabolize sucrose.

**Figure 2.**
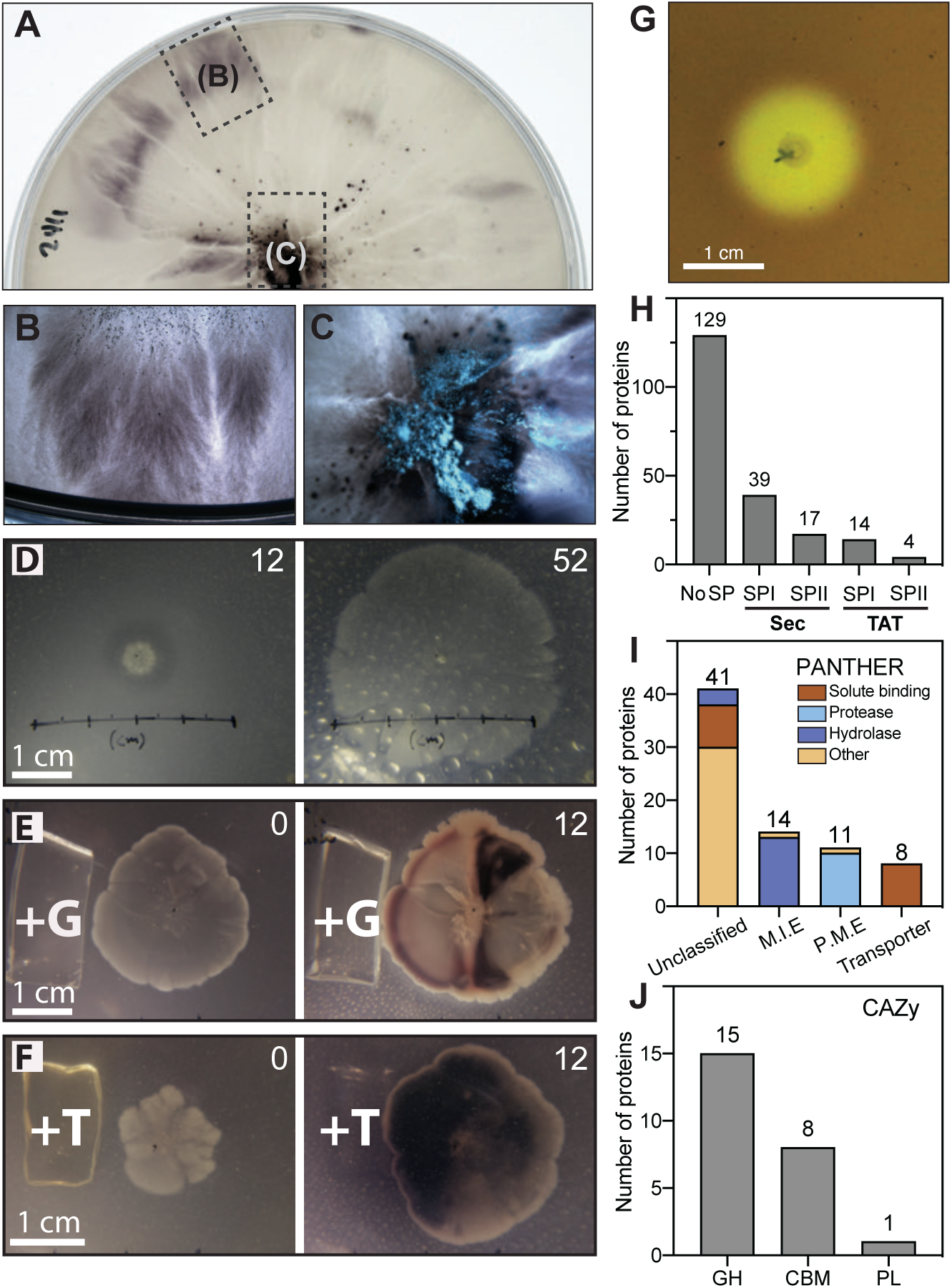
Nutrient poor agar media promote the *S. coelicolor* foraging phenotype. (**A**) *S. coelicolor* spores in glycerol storage buffer inoculated on FAM. The foraging phenotype covering the petri dish (⌀ 8.8 cm) was photographed after 15 weeks of incubation at 30°C. Transparent pigment-free and lightly pigmented sectors are visible. (**B**) A photograph of a pigmented sector extending towards the edge of the dish. (**C**) Formation of aerial hyphae were only observed at the site of spore inoculation. (**D**) Time-lapse stills showing the foraging growth progression at day 12 and 52. (**E**) Time-lapse stills after addition of glucose (+G) to a foraging colony at day 0 and 12. (**F**) Time-lapse stills of addition of tryptone (+T) to a foraging colony at day 0 and 12. (**G**) Photograph of foraging *S. coelicolor* grown on FAM, stained with Lugol’s solution. (**H**) Number of proteins containing a signal sequence in the foraging secretome, predicted using SignalP 6.0 [30]. Categories include general secretory (Sec) substrates cleaved by SPase I (Sec/SPI) or SPase II (Sec/SPII, lipoproteins) and twin-arginine substrates cleaved by SPase I (Tat/SPI) or SPase II (Tat/SPII, lipoprotein), and “No SP” for no predicted signal peptide. (**I**) PANTHER functional classification of proteins from the foraging secretome with predicted signal peptides. The graph shows the number of proteins assigned to each protein class. (**J**) CAZy classification of the foraging secretome: Glycoside Hydrolases (GH), Polysaccharide Lyases (PLs) and Carbohydrate-Binding Modules (CBMs).

The progressive expansion and sector formation of foraging colonies on FAM were studied using time-lapse imaging (Figure 2D, Video S2). Introduction of glucose or tryptone to a foraging colony induced raised colony growth and strong pigment production, typical for growth in nutrient-rich conditions (Figure 2E, 2F, Video S3 and Video S4).

Agar was utilized as the main carbon source when growing on a deprived media such as FAM, and the agar degradation was visualized by staining the plates with iodine-based Lugol’s solution which revealed the distinct halo around the colonies indicating substrate degradation (Figure 2G). Proteins were extracted from the zones outside of foraging colonies and analyzed using mass spectrometry. Signal peptide prediction using SignalP 6.0 [30] was performed on all *S. coelicolor* coding sequences, identifying 696 proteins with predicted signal peptides. Among the 203 proteins identified from the foraging secretome, 56 were predicted to have a Sec translocon signal peptide, 18 had a Tat translocon signal peptide, and 123 lacked any signal peptide (Figure 2H, Table S1). PANTHER classification of signal peptide-containing proteins categorized them into four categories (Figure 2, Table S2). The metabolite interconversion category featured 13 hydrolases and one oxidoreductase, while the protein modifying enzyme category included ten proteases and a ubiquitin-protein ligase. Among the unclassified proteins were additional hydrolases and solute binding proteins. The transporter category included the ABC transporter BldKB, known for its role in carbon catabolite repression and mycelium differentiation [31, 32]. Four proteins in the foraging secretome, encoded by genes containing the *bldA*-specific TTA codon, demonstrated that *bldA* is active during foraging growth: an uncharacterized protein without a signal peptide (Q9L2F9, SCO2524), a carbohydrate-binding module (CBM) domain-containing hydrolase (Q9RKF0, SCO3487) with a Sec-signal peptide, and two uncharacterized secreted proteins with Sec-signal peptides (Q9RKE0, SCO3498 and Q9K452, SCO7233) (Table S3). However, a *bldA* mutant still displayed the foraging phenotype, suggesting that BldA is expressed but not essential for the phenotype. Mutants of the classical Whi-regulator proteins, that act downstream of BldA in the regulatory cascade also displayed the foraging phenotype (Figure S2B). In total, 24 secretome proteins associated with hydrolytic and non-hydrolytic cleavage of glycosidic bond and carbohydrate adhesion were identified in the Carbohydrate-Active enZYmes (CAZy) Database [33] (Figure 2J).

### Distinct behavioral differences between foraging and conventional phenotypes

Colonies grown on FAM and TSA were imaged in high resolution using scanning electron microscopy (SEM) imaging to clearly visualize the two types of growths. First, the predominantly submerged growth in FAM was confirmed.

Streptomyces growth on TSA showed a consistently thick layer of hyphae piled on top of each other (Figure 3A, B). The piled-up surface growth continued to the outskirts of the colonies (Figure 3C). In contrast, few hyphae were seen at the surface on FAM except a few displaying a “dolphin-like” behavior (Figure 3D), with the exception of small patches of hyphae at the site of inoculation and occasionally at other sites on the surface. Cross sections of colonies confirmed that nutrient-rich growth resulted in limited number of hyphae inside the agar, while the majority of growth occurred inside the agar on FAM (Figure 3E, F), with numerous of clearly distinguishable hyphae within the colony in contrast to the agar material alone (Figure 3G, H). The high-resolution view demonstrates that, in an environment with low levels of low-molecular-weight organic compounds and abundant polysaccharides, the foraging colony infiltrates the substrate.

**Figure 3.**
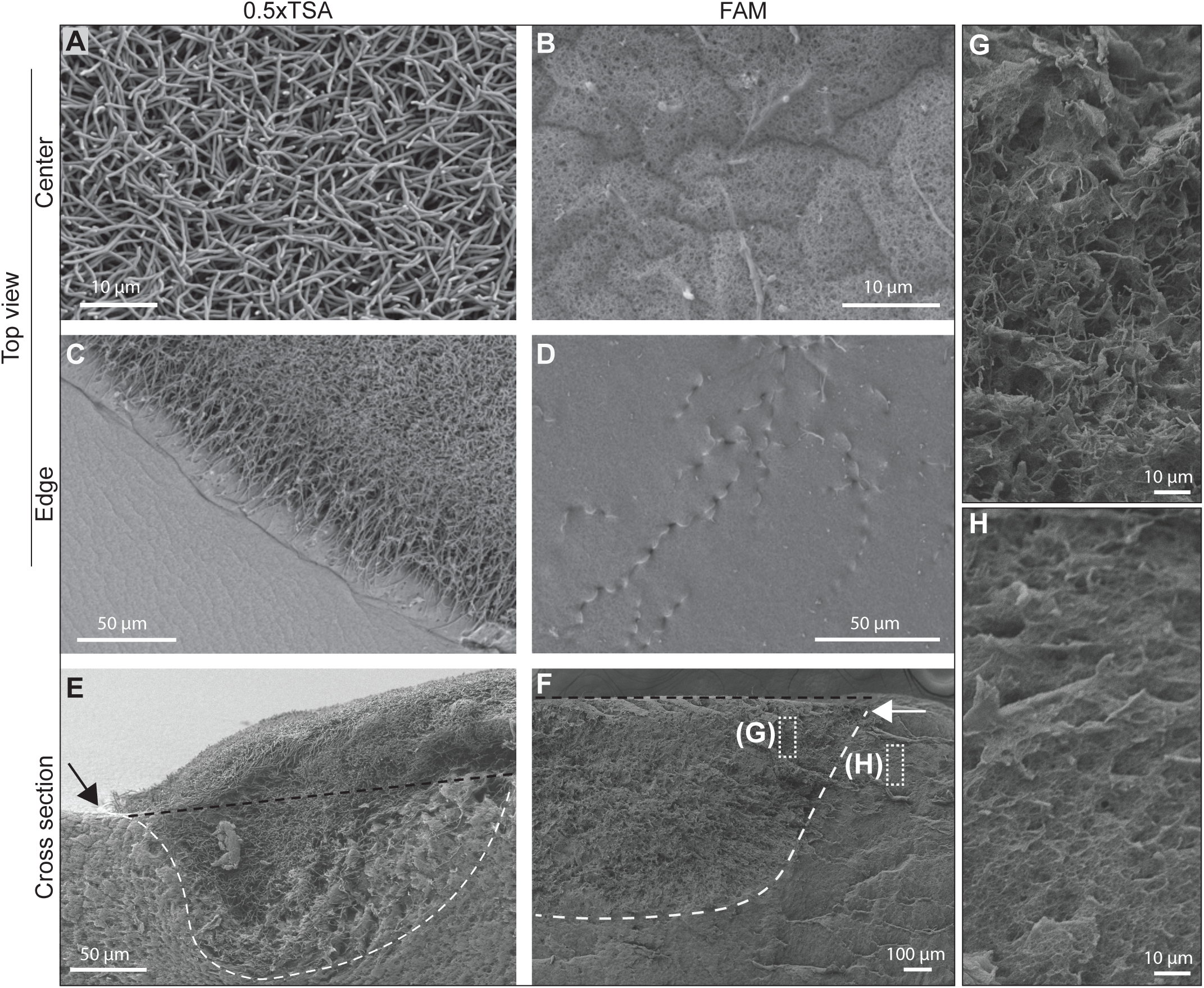
Foraging hyphae are primarily found in the agar substrate. SEM imaging of *S. coelicolor* grown on FAM and 0.5xTSA for 7 days. The glycerol storage buffer was washed away and replaced with water. (**A**) Top view of the center of a colony grown on TSA. (**B**) Top view of the center of a foraging colony. (**C**) Top view of the edge of a TSA colony. (**D**) Top view of the edge of a foraging colony shows few hyphae at the surface of the FAM. (**E**) Cross section of a TSA-colony show hyphae predominantly grow on top on 0.5xTSA. (**F**) Cross section of a foraging colony shows that hyphae extending through the agar are mainly observed on the FAM-plates. Black dotted lines indicate the agar surface in the cross sections. White dotted lines encompass where hyphae are found. (**G**) A magnified view shows hyphae inside the FAM. (**H**) A magnified view of FAM showing absence of hyphae outside of the foraging colony.

To further investigate potential differences between growth at different nutrient availability, we turned the attention to the sectors formed during foraging growth (Figure 1A, 2A, 2B, Video S1 and S2). Since significant genomic instability in *S. coelicolor* colonies grown on rich media has been reported [13, 14], we isolated and sequenced foraging sectors to investigate whether they are formed as a result from this genomic instability. A lineage from the initial inoculation, a subsequent sector ‘A’, and its subsequent sectors ‘I’ and ‘IV’, were sequenced using the Illumina platform (Figure S3A). Analysis using breseq [34] showed no evidence of terminal deletions in foraging sectors. Point-mutations were found in three genes but no influence on growth were detected in either rich or nutrient-deprived conditions (Figure S3A). These results emphasize that the adaptation strategies significantly differ between nutrient-rich and nutrient-deprived environments. The absence of major genomic deletions on FAM suggests an importance to maintain the integrity of the genome for survival in nutrient limited conditions.

### Foraging phenotype is found across the phylogeny of *Streptomyces*

To assess whether foraging growth was unique to *S. coelicolor*, the study was extended to include other *Streptomyces* species representing different branches of the phylogenetic tree, as well as a species from the closely related genus *Kitasatospora*. All 12 strains tested displayed the foraging growth phenotype on FAM (Figure 4A, B), and agar degradation was confirmed by Lugol’s solution staining (Figure S3B). Similar to *S. coelicolor*, foraging growth was inhibited in all strains with the addition of 1% glucose or tryptone to FAM (Figure S3C, D).

**Figure 4.**
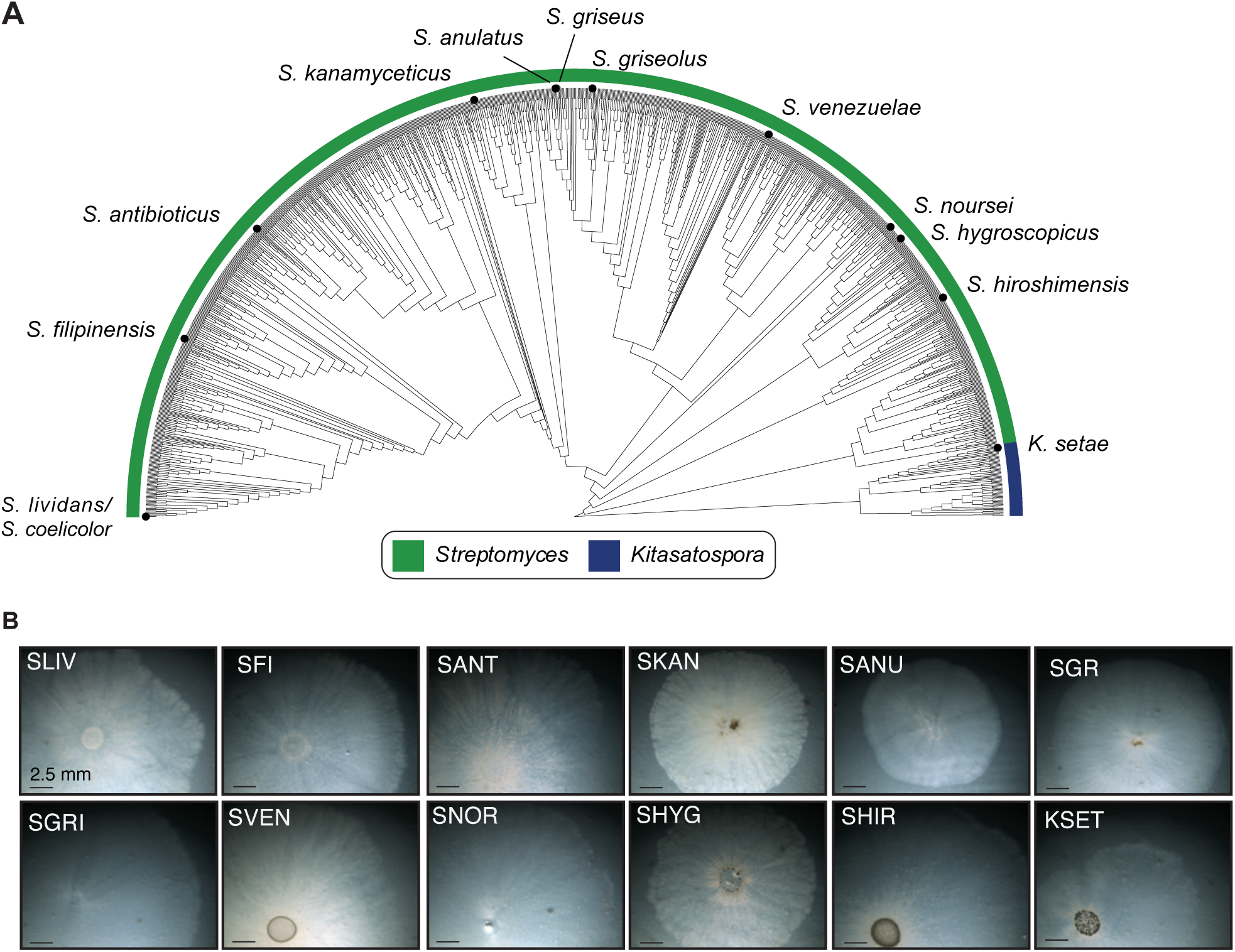
Phylogenetic tree of the Streptomyces and Kitasatospora genera. (**A**) The phylogenetic tree, displayed as a half-circle, was generated using GTDB-Tk and pruned to include only the *Streptomyces* (blue) and *Kitasatospora* (green) genera. The positions of the species tested for foraging phenotype are indicated with a point on the branch and the species name. Species confirmed to exhibit the foraging phenotype are indicated by points on their respective branches, along with their NCBI species names. (**B**) Microscopy images showing the foraging phenotype of each *Streptomyces spp.* cultivated for three weeks on FAM on the right side of the tree. Abbreviations: SANT; *S. antibioticus*, SANU; *S. anulatus*, SCO; *S. coelicolor*, SFI; *S. filipinensis*, SGR; *S. griseus*, SGRI; *S. griseolus*, SHIR; *S. hiroshimensis*, SHYG; *S. hygroscopicus*, SKAN; *S. kanamyceticus*, SLIV; *S. lividans*, SNOR; *S. norsei*, SVEN; *S. venezuelae*, KSET; *Kitasatospora setae*.

To investigate whether foraging was a characteristic that could be used to distinguish a streptomycetes, local soil samples were incubated on FAM, and colonies displaying the foraging phenotype were analyzed using MALDI-TOF to determine the species. A colony displaying the foraging phenotype was picked at random from each of the seven soil samples. All seven colonies were confirmed to be streptomycetes (Table S4). These findings further strengthened the ubiquity of foraging growth across the *Streptomyces* phylogeny. Until recently, it was uncertain whether *Kitasatospora* was a separate genus from *Streptomyces but* has been elucidated since [35–37]. The fact that *K. setae* also displayed foraging demonstrates that the phenotype is not confined to the *Streptomyces* genus but also occurs in at least one closely related genus.

### Foraging streptomycetes demonstrates increased antibacterial activity

Since streptomycetes are known for their production of antimicrobial metabolites and also encode numerous BGCs that are not expressed during conventional growth, the potential of foraging colonies to inhibit bacterial growth was investigated. Due to the nutrient-depleted state of FAM, *B. subtilis* and *E. coli* were unable to proliferate directly on the agar surface. Therefore, to test for antibacterial activity, FAM with foraging *S. coelicolor* colonies was overlaid with diluted lysogeny agar (LA) and *B. subtilis* 168 and *E. coli* MG1655 then added on top of the LA surface. The growth was evaluated after incubation using a stereo microscope because of the smaller colonies formed due to the slower proliferation of the indicator bacteria on the diluted LA. This method was applied to the 12 additional *Streptomyces* species to also assess their antibacterial properties during foraging. Most species showed increased capacity to inhibit both indicator bacteria on nutrient-depleted FAM (11/13) compared to FAM supplemented with glucose (FAM+G, 3/13) and tryptone (FAM+T, 1/13) (Figure 5A). However, *S. coelicolor* did not exhibit enhanced antibacterial activity on FAM compared to the nutrient-supplemented media. Performing the same overlay assay on *S. coelicolor* developmental mutants showed that the *bldA* mutant lost its ability to inhibit *B. subtilis* growth, whereas the *adpA* mutant acquired the ability to inhibit *E. coli* (Figure 5B).

**Figure 5.** Metabolome and antimicrobial activity of foraging colonies. (**A**) Bactericidal activity of foraging streptomycetes determined by overlay-assays with indicator bacteria (Ec=*E. coli*, Bs=*Bacillus subtilis*) on FAM, FAM+G and FAM+T. (**B**) Bactericidal activity of *S. coelicolor* developmental mutants as determined by overlay assays on FAM. Dark purple indicates inhibition of indicator bacteria, light purple indicates partial growth inhibition, and grey indicates that there was no detectable effect. (**C**) Inhibition zones around *S. coelicolor* on FAM contaminated by *P. citrinum* mold. (**D**) Inhibition zones around *S. coelicolor* wild-type (WT), bldA and M851 (*adpA*-mutant) inoculated with *P. citrinum*. (**E**) Scores plot of LC-MS analysis in negative detection mode of FAM and FAM+G-agar inoculated with *S. coelicolor*. (**F**) Loadings plot from negative detection mode. Features with relatively elevated levels on FAM are highlighted in blue and features with relatively elevated levels in FAM+G-agar are highlighted in red. (**G**) Scores plot of LC-MS analysis in positive detection mode of FAM and FAM+G-agar inoculated with *S. coelicolor*. (**H**) Loadings plot from positive detection mode. Features with relatively elevated levels on FAM-agar are highlighted in blue and features with relatively elevated levels in FAM+G-agar are highlighted in red.

*S. coelicolor* colonies grown on FAM were found to inhibit the growth of a contaminating mold (Figure 5C), identified as *Penicillium citrinum* by 18S/ITS sequencing. This inhibition was replicated by growing *S. coelicolor* on FAM for five days before inoculating *P. citrinum* spores at a distance of 2 cm, resulting in an average 4.3 mm inhibition zone (±0.76 mm, n = 96). Inhibition was not observed when glucose was added to the agar or on any other nutrient-rich media, nor on FAM with pH below 7. Interestingly, the *bldA* mutant, typically unable to produce certain antimicrobial metabolites, inhibited *P. citrinum* growth on FAM. However, the *adpA*-mutant strain M851 showed reduced inhibition compared to the wild-type (Figure 5D). All tested *whi*-mutants *(-A, -B, -G, -H,* and *-I*) exhibited mold-inhibitory activity (Figure S4A). Inhibition of a local *Aspergillus spp.* isolate was also demonstrated using an overlay assay with a 100x diluted TSA-overlay, since the *Aspergillus spp.* did not grow on FAM alone (Figure S4B). However, no inhibition of *Saccharomyces cerevisiae* or *Candida albicans* was detected using the same assay. To our knowledge, there are no reports of antifungal activity for *S. coelicolor* A3(2), despite it being extensively studied for its pigment and metabolite production since the 1950s [38]. To investigate whether mold inhibition was a common property during foraging, the 12 additional *Streptomyces* species were tested for mold inhibition, but the mold inhibitory activity was specific to *S. coelicolor*, as none of the other species inhibited *P. citrinum*.

The altered antimicrobial activates suggest that the foraging characteristics extend beyond genomic stability and colony morphology and include significant changes to metabolite production.

### Foraging *S. coelicolor* displays an altered metabolomic profile

To determine whether there were any differences in the metabolomic profiles of *S. coelicolor* grown on FAM and FAM supplemented with 1% glucose (FAM+G), agar was sampled around the colonies for untargeted mass spectrometric analysis. The metabolomic profiles differed between FAM and FAM+G, both in negative (anions, goodness of fit: R2(X)=0.75, goodness of prediction: Q2(X)=0.97) and positive detection modes (cations, R2(X)=0.63, Q2(X)=0.47) (Figure 5E, F). Some features with relatively higher concentrations on FAM-agar (highlighted in blue in Figure 5G, H) could be matched by monoisotopic molecular weight with known *S. coelicolor* metabolites from the StreptomeDB [39]. Among these annotated features, the signaling molecule γ-butyrolactone was found at higher concentrations in FAM+G. Wailupemycin G was the only annotated metabolite found at considerably higher concentrations in FAM (Figure S4C, Table S5). There were an additional 30 features, with higher levels in the FAM condition, that could not be annotated using their monoisotopic mass and StreptomeDB (Figure S4D, E, Table S6). Overall, more metabolites features were detected in the FAM+G samples. Twelve of these features were unique to FAM and were thus either present below detection limits or not produced on FAM+G. Genetic analysis of BGCs has revealed that each *Streptomyces* species could have the capacity to produce over 20 secondary metabolites [17, 21–23]. However, standard cultivation often fails to stimulate the activation of the regulatory pathways responsible for their synthesis [24, 25]. The mass spectrometry results demonstrate a significant alteration in the metabolomic profile of foraging *S. coelicolor* as a consequence of the different environmental stimuli.

The gained ability of *S. coelicolor* to inhibit mold-growth together with the overall enhanced antibacterial activities of other *Streptomyces* species suggests that there are also alterations of their metabolomes. This shows potential for using FAM growth in screening for active secondary metabolites not produced on rich media.

## Discussion

The primary habitat of streptomycetes, the soil, is a dynamic environment with many variable environmental factors. Traditional laboratory cultivation using a limited number of rich media is unlikely to represent the full repertoire of adaptations in a diverse and competitive environment such as the soil. In this study, we demonstrate that global nutrient depletion in growth media coincides with a morphological transition of *S. coelicolor* colonies to a previously undescribed phenotype. We chose the term “foraging” to describe this phenotype because the colony demonstrates continuous outward expansion on nutrient-depleted substrates rich in agar polysaccharides, a process that is halted when a low–molecular-weight carbon source (e.g., glucose) is introduced. This behavior mirrors a forager’s search for and subsequent exploitation of available resources.

Media composition is known to be crucial for *Streptomyces* life-cycle progression, with nutrient depletion often referred to as one of the triggers of differentiation and sporulation, though the process is acknowledged to be complex [2, 40–42]. Observations of *S. coelicolor* colonies over time, combined with metabolomics analysis, showed that the transition to the foraging phenotype on 0.5xTSA coincided with nutrient depletion and not differentiation and subsequent sporulation. This demonstrates that sporulation under nutrient-depletion is conditional and that supplementation of certain polysaccharides can also induce sporulation. Furthermore, it underscores that the use of conventional rich media has not revealed the full repertoire of *S. coelicolor* adaptive regulation. This ability to continue growth in a nutrient-poor environment allows *S. coelicolor* to avoid entering a dormant state as spores awaiting a rich supply of nutrients, enabling continuous growth. It was recently demonstrated that secondary metabolite production is significantly increased as a result from genetic instability on nutrient rich agar [13]. However, we could not find any support for major mutational deletions under nutrient limited foraging growth when investigating the sectors formed during foraging growth. This further emphasizes that the survival strategies of *S. coelicolor* differ between nutrient-depleted and nutrient-rich agar growth.

Fluctuations in metabolite levels during growth results from the breakdown and uptake of external sources and the subsequent release into the surroundings [43]. Four distinct trends in metabolite dynamics were distinguishable throughout the *S. coelicolor* transition to foraging: depletion, accumulation, initial accumulation followed by depletion, and unchanged levels. The trend of initial accumulation followed by depletion is likely due to the breakdown of macromolecules from the TSA media and subsequent consumption. Accumulation of metabolites may occur due to by-product overflow or overflow-independent mechanisms, such as strong transport affinity [44]. Supplementation of the accumulated metabolites to media did not trigger foraging growth, demonstrating that their accumulation is not promoting the phenotype. Instead, the release of metabolites might be a deliberate strategy to interact with other organisms in the natural habitat of *S. coelicolor*. Two accumulated metabolites, β-alanine and pantothenic acid, are crucial in various biochemical pathways. β-alanine, essential for pantothenic acid synthesis – a precursor of Coenzyme A – enhances respiration in alkaline soils [45].

In contrast to rich media, where low-molecular-weight organic compounds are abundant [29, 46], soil environments typically have transient availability of such nutrients [47, 48]. Fertile soils are rich in complex carbohydrates and plant debris and have a flux of available nutrients throughout the year to which soil inhabitants must adapt [49]. FAM, our minimal defined agar medium, where agar acts as the sole carbon source, supports spore germination and foraging growth, but the addition of easily metabolized nutrient (e.g., glucose) promotes conventional colony phenotypes and sporulation. This indicates that the complex polysaccharides in agar are not preferred over simpler organic compounds. Nitrogen and phosphate also play a crucial role in cell cycle progression and metabolite production. The FAM media used here, with higher nitrogen levels than typically found in soils [47], may influence the phenotype. Elevated levels of nitrogen in rich media have been associated with repression of secondary metabolite production [50]. Furthermore, elevated levels of inorganic phosphate have been shown to inhibit the production of several secondary metabolites [51–53], and there has been shown that there is a significant flux of organic nitrogen in soils using microdiffusion experiments [54–56]. Supplementation of FAM with an organic nitrogen source (i.e., asparagine), inhibited foraging growth and promoted conventional colony growth. Therefore, the levels and types of phosphate and nitrogen sources are also contributing to the foraging phenotype on FAM.

The prospect of finding new metabolites produced by novel colony morphologies has been previously demonstrated for the exploratory phenotype that certain streptomycetes exhibit when co-incubated with yeast or when grown on rich media with specific media components alone [20, 57]. Inhibition of molds by foraging *S. coelicolor* A3(2) is, to our knowledge, the first such observation in one of the most extensively studied *Streptomyces* species [58]. This underscores the novelty of the metabolic changes induced by foraging, given that nutrient-rich substrates commonly used for cultivation apparently do not provide the environmental stimuli required for producing the active molecule. The mold was identified as a *Penicillium* species, but inhibition was also observed against an *Aspergillus* species. By contrast, no inhibition occurred with the yeast *S. cerevisiae* or the dimorphic fungus *C. albicans* (capable of both single-cell and filamentous growth), suggesting a specificity of the inhibition. A mold-specific activity could be advantageous for antifungal drug development, as it may pose less toxicity toward mammalian cells, a major challenge in current antifungal therapies [59]. However, further research is needed to characterize the precise range of susceptible species, isolate the active compound(s), and elucidate the mechanism by which mold growth is inhibited. The gained ability of foraging *S. coelicolor* to inhibit mold-growth, together with the overall enhanced potential of other *Streptomyces* species to inhibit Gram-positive and Gram-negative bacteria, underscores a competitive advantage for foraging colonies in nutrient-deprived conditions and suggests that the altered metabolomic profile detected for *S. coelicolor* might be a general phenomenon. The low nutrient levels used for the overlay assays, to avoid interfering with the foraging colonies, will naturally also influence the target bacteria rendering them either more or less susceptible to the antimicrobial effects of foraging colonies [60–62], a factor which should be considered in future studies. It is well established that growth conditions, including the choice of carbon source, is one among strategies used to influence secondary metabolism and can activate silent BGCs in streptomycetes [63, 64] Altogether, there is potential for using FAM growth in screening for biologically active secondary metabolites not produced on rich media.

The foraging *S. coelicolor* secretome contained 74 proteins with predicted signal peptides. Despite the presence of BldA-dependent proteins in the foraging secretome, a *bldA* mutant still displayed foraging growth, demonstrating that *bldA* regulation is not essential for this phenotype. The regulatory mechanism of BldA-dependent *adpA* translation [6] and the AdpA regulation of *bldA* [12] in combination with the activation of downstream regulatory genes by both factors appear to be of importance for the antimicrobial activity of the foraging colonies. The finding that a foraging *ΔbldA* mutant loses the ability to inhibit *B. subtilis*, whereas a foraging *ΔadpA* mutant gains the ability to inhibit *E. coli*, suggests that inhibition of *B. subtilis* by *S. coelicolor* is positively regulated by *bldA*, while *adpA* negatively controls inhibition of *E. coli*, directly or indirectly. Conversely, mold inhibition appears to be positively regulated by *adpA,* as the *ΔadpA* mutant showed no inhibition zones. Therefore, although *bldA* is not required for the foraging colony morphology nor mold inhibition, it is required for the inhibition of *B. subtilis*. These observations highlight the complex regulatory roles of *bldA* and *adpA* in antimicrobial regulating the antimicrobial activities during foraging growth.

Unlike exploratory growth, which so far has only been observed in some species [20], foraging growth was observed in all tested *Streptomyces* species from different parts of the phylogeny, indicating it may be a common feature across the genus. Further, *K. setae* from the closely related genus *Kitasatospora* also displayed the foraging phenotype, demonstrating that the foraging is not limited to streptomycetes. The mechanism by which foraging growth is regulated remains to be elucidated. Identifying the regulatory components would result in understanding the required nutrient deprived conditions. Importantly, understanding its regulation could be used to genetically activate the production of certain secondary metabolites.

In summary, this study demonstrates a previously overlooked growth phenotype of *Streptomyces* with distinctly different colony morphology and metabolite profiles compared to conventional growth on rich media. The nutrient-deprived growth has a distinct phenotype and an altered metabolome, with enhanced antimicrobial properties. The “problem of rediscovery” of known antimicrobial compounds [65], despite the presence of numerous silent BGCs, is a significant challenge to overcome in combating antimicrobial resistance. This novel growth behavior offers new insights into this issue and paves the way for discovering new regulatory mechanisms underlying the foraging growth phenotype. Taken together, this novel growth phenotype provides a new perspective on the environmental adaptations and competitiveness of streptomycetes under nutrient-deprivation, with potential implications for antimicrobial drug discovery.

## METHODS

### Strains, media, and culture conditions

The strains used in this study are listed in Table S7. *Streptomyces* strains were grown on foraging agar medium (FAM) (1 g (NH_4_)_2_SO_4_, 0.5 g K_2_PO_4_x7H_2_O, 0.1 g FeSO_4_x7H_2_O, 10 g agar in 1L pH 7.5) [29]. FAM were supplemented with 1% of various sugars: glucose (FAM+G), sucrose (FAM+S), fructose (FAM+F) or tryptone as an amino acid source (FAM+T) when indicated. To investigate the effect of pH on mold-inhibition the pH of FAM was set using HEPES-buffer and HCl/NaOH. Tryptone soy broth (Thermo Fisher Scientific) was prepared according to instructions and diluted 0.5x when indicated with water and 1.5% agar added before autoclaving. MS-agar (20 g mannitol, 20 g Soy flour, 15 g agar in 1L) agar were used for sporulation and petri dish time-lapse imaging. Effects of accumulated metabolites were conducted by adding 20 mg/ml: beta-alanine, meso-erythriol (Merck), Potassium-citramalate monohydrate (Merck), urea (Merck), and pantothenic acid (Merck). All incubations were carried out at 30°C with a vessel containing water to maintain high humidity and minimize plate drying. Plates were sealed with Parafilm and regularly inspected; seals were replaced as needed to prevent drying during prolonged incubation.

### Isolation and identification of local isolates

The inhibited mold was isolated from the FAM dish and recultivated on TSA. The agar plate was sent to Eurofins for 18S/ITS sequencing. For isolation of local streptomycetes, soil samples from potatoes and plant soil were dried at room temperature before being deposited on FAM. The plates were incubated at 37°C until foraging colonies were observed after approx. 5–10 days. Colonies with characteristic foraging phenotypes were re-streaked on TSA and characterized using MALDI-TOF (Bruker). The local Aspergillus spp. was isolated from a soil sample and assessed morphologically.

### Colony imaging

For all imaging of small colonies spores were inoculated on agar media and visualized using a Nikon SMZ1500 dissection microscope and a Nikon Digital sight DS-Fi1 camera, with either surface illumination using LEDs or oblique sub-stage illumination. Whole petri dishes were imagined using a Nikon D7000 DSLR. For time-lapse imaging, the lids of each plate were washed with detergent, dried, and then wiped with 10% Triton X-100 to reduce droplet formation. Plates were carefully sealed with Parafilm to prevent drying during the time-lapse imaging. Agar time-lapses were recorded using a Raspberry Pi 3 model B with camera module v2 mounted in a 10x lens (Olympus). The device was placed in a 30°C dark room with an LED lamp programmed to illuminate the plate only during image capture, one frame per hour. ImageJ distribution Fiji [66] was used to process single frames into videos and Adobe Photoshop CC software for still images.

### Electron microscopy

Colonies of *S. coelicolor* on agar were cut, fixed overnight at 4°C in 2.5% glutaraldehyde/0.1M sodium cacodylate buffer, dehydrated in series of ethanol, critical point dried and coated with a 5 nm Au/Pd before being analyzed with a Merlin Field Emission SEM (Zeiss) using SE-HE2 and in-lens detectors. Images were processed using Adobe Photoshop CC software.

### Inhibition assays

For mold inhibition, *Streptomyces* spores were inoculated on FAM and incubated at 30°C for 5 days. Spores from *P. citrinum* were inoculated 2 cm from the foraging colony. The plates were incubated for 14 days and observed daily for inhibition zones and imaged as described above. For bacterial inhibition, plates with *Streptomyces* spp. were incubated for 20 days before a thin layer of 100x diluted TSA (1.5% agar) was placed on top of agar and incubated overnight at 4°C. Overnight cultures of *E. coli* and *B. subtilis* were diluted and 2×10^7^ CFU in 100 µl was added to the agar overlays and incubated overnight at 37°C. Inhibition zones were assessed and imaged using a Nikon SMZ1500 dissection microscope and a Nikon Digital sight DS-Fi1 camera.

### Foraging secretome protein analysis

The agar outside of foraging *S. coelicolor* on FAM incubated for 20 days was extracted. The agar was crushed and incubated overnight with water in a 50 ml Falcon tube. Proteins in the supernatant were precipitated by addition of TCA and incubated on ice for 4 minutes before centrifugation at 20,000g for 5 min. The pellet was washed twice with 200 μl ice-cold acetone. The pellet was dried, dissolved in protein sample buffer and loaded on an SDS-PAGE. Protein dense regions were cut and sent for mass spectroscopy identification to Linköping University, where general in-gel digestion was performed as described by Shevchenko et al. (Shevchenko, Tomas, Havlis, Olsen, & Mann, 2006).

### Metabolomic analysis

Metabolic profiling of media nutrients by GC-MS was performed at the Swedish Metabolomics Center in Umeå, Sweden. For the nutrient analysis of agar, six replicates of agar plugs were collected from the edge of the dish and 0.5 cm away from the *S. coelicolor* colonies on 0.5xTSA medium. This sampling was performed on six different ⌀ 4 cm plates. Three control samples were taken at every time point from three sterile 0.5xTSA dishes. Sample preparation was performed according to A et al. (A et al., 2005). The samples were stored at −80 °C until analysis. Small aliquots of the remaining supernatants were pooled and used to create quality control (QC) samples. The samples were analyzed in batches according to a randomized run order on GC-MS. Derivatization and GCMS analysis were performed as described previously A et al. (A et al., 2005). Non-processed MS-files from the metabolic analysis were exported from the ChromaTOF software in NetCDF format to MATLAB-R2020a (Mathworks, Natick, MA, USA), where all data pre-treatment procedures (base-line correction, chromatogram alignment, data compression and Multivariate Curve Resolution) were performed. The extracted mass spectra were identified by comparisons of their retention index and mass spectra with libraries of retention time indices and mass spectra (Schauer et al., 2005). Mass spectra and retention index comparison was performed using NIST MS 2.2 software. Annotation of mass spectra was based on reverse and forward searches in the library. Masses and ratio between masses indicative of a derivatized metabolite were especially notified. The mass spectrum with the highest probability indicative of a metabolite and the retention index between the sample and library for the suggested metabolite was ± 5 the deconvoluted “peak” was annotated as an identification of a metabolite. The annotated nutrients were plotted using GraphPad Prism (version 9.0.2) to show trends over time and ridge plots were generated using R version 4.2 and packages ggplot2 3.4.4 and ggridges 0.5.4.

For untargeted metabolite analysis, agar stubs were punched out from just outside of *S. coelicolor* colonies on FAM and FAM+G. For each condition, twelve samples were isolated from plates inoculated with *S. coelicolor*, in addition to two control samples each from sterile plates. The weight of the samples was adjusted to 100 mg before the addition of 1 mL extraction buffer (90/10 v/v methanol:water). Internal standards (13C9-Phenylalanine, 13C3-Caffeine was, D4-Cholic acid, D8-Arachidonic Acid, 13C9-Caffeic Acid) were added to each sample. The sample was shaken at 30 Hz for 3 min in a mixer mill, and proteins were precipitated at +4 °C for 2h on ice. The sample was centrifuged at +4 °C, 14 000 rpm, for 10 min. The supernatant was transferred to a microvial and solvents evaporated. Before analysis, the sample was re-suspended in 10 + 10 µL methanol and water. The set of samples were analyzed in positive and negative mode. The chromatographic separation was performed on an Agilent 1290 Infinity UHPLC-system (Agilent Technologies, Waldbronn, Germany). Re-suspended aliquots (2 µL) of the agar extracts were injected onto an Acquity UPLC HSS T3, 2.1 x 50 mm, 1.8 µm C18 column in combination with a 2.1 mm x 5 mm, 1.8 µm VanGuard precolumn (Waters Corporation, Milford, MA, USA) held at 40°C. The gradient elution buffers were A (H2O, 0.1 % formic acid) and B (75/25 acetonitrile:2-propanol, 0.1 % formic acid). The compounds were detected with an Agilent 6550 Q-TOF mass spectrometer equipped with a jet stream electrospray ion source. The settings were kept identical between the modes, except the capillary voltage. The data processing was performed using the Recursive Feature Extraction algorithm within Agilent Masshunter Profinder version B.08.00 (Agilent Technologies Inc., Santa Clara, CA, USA). All multivariate statistical investigations (PCA) were performed using the software package SIMCA®-P+ version 15.0.2 (Umetrics, Umeå, Sweden). Annotation of unknown metabolites were performed by comparing monoisotopic molecular weights with *S. coelicolor* metabolites in the StreptomeDB (Moumbock et al., 2021) using R software (4.2). Metabolomic features were plotted using GraphPad Prism (version 9.0.2). All raw mass spectrometry files used in this study have been deposited to ScieLifeLab Data Repository with the dataset identifier DOI: https://doi.org/10.17044/SCILIFELAB.25593480 and https://doi.org/10.17044/SCILIFELAB.25593528.

### Phylogenetic analysis

Phylogenetic analysis was conducted to investigate the evolutionary relationships among *Streptomyces* species exhibiting the foraging phenotype. Phylogenomic data were obtained from the Genome Taxonomy Database Toolkit (GTDB-Tk) release 220.0, accessed in April 2024. The phylogenetic tree file (bac120.tree) and associated taxonomy and metadata files (bac120_taxonomy.tsv and bac120_metadata.tsv) were imported into R version 4.2 and processed using the ape 5.8, tidyverse 2.0.0, readr 2.1.5, and ggtree 3.12.0 packages [67–70]. Taxonomic data were processed to filter species belonging to the genus *Streptomyces* and the species *Kitasatospora*. To ensure non-redundancy, only one representative accession per species was included by selecting distinct species entries from the dataset. The phylogenetic tree was pruned to include only the selected genus using the “drop.tip” function from the ape package. The final pruned tree was visualized using the ggtree package [67]. The positions of each species tested for foraging were annotated to visualize the phylogenetic relationships among the selected species.

### DNA sequencing of foraging sectors

Two levels of sectors formed during foraging on FAM were grown in TSB-media and 600 mg of pellet was sent to MicrobesNG for Illumina sequencing. Quality controlled and trimmed reads with 30x coverage were obtained and analyzed using breseq software [34] with default settings, using the *S. coelicolor* A(3)2 reference genome (Genbank: GCA_000203835.1, [17]) updated using wild-type sequence data to accommodate for genomic variation in lab strain. The reads of descendent sectors were mapped against this updated reference genome. Illumina read data used in this study have been deposited to SciLifeLab Data Repository with the dataset identifier DOI: https://doi.org/10.17044/SCILIFELAB.25593405.

### Statistical analysis

In this study, certain measurements in the untargeted metabolomics analysis fell below the detection limits, resulting in missing values. To ensure an accurate representation of our findings and maintain the integrity of the statistical analysis, these missing values were omitted from the dataset. Consequently, all values presented in the violin plots depict observable data trends and variations above the detection threshold. All statistical analyses were conducted using GraphPad Prism (version 9.0.2). Details on the specific type of statistical analysis and the criteria for significance testing are provided in the legends of the respective figures.

## Acknowledgment

We thank National Microscopy Infrastructure, NMI (VR-RFI 2016-00968) at Umeå Centre for Electron Microscopy (UCEM). Swedish Metabolomics Centre, Umeå, Sweden (www.swedishmetabolomicscentre.se) is acknowledged for metabolic profiling by LC-MS and GC-MS. Illumina sequencing was provided by MicrobesNG (http://www.microbesng.com). Postdoc fellowship funding from the Kempe foundations and Sven och Lilly Lawskis foundation. Project granted by Umeå Centre for Microbial research (UCMR) and Swedish Foundation for Strategic Research (RIF21-0067). Anders Sjöstedt for MALDI-TOF analysis of streptomycetes and Ala Javadi for sample preparation. We are grateful to Cecilia Drakskog for her thorough and constructive feedback on the manuscript.

## Resource availability

### Materials availability

This study did not generate new unique reagents.

### Data and code availability

The Illumina sequencing data, the metabolomics and nutrient mass spectrometry data has been deposited in the SciLifeLab Data Repository (Sandblad, Linda; Söderholm, Niklas, 2024) under DOI: https://doi.org/10.17044/SCILIFELAB.25593528, https://doi.org/10.17044/SCILIFELAB.25593480, https://doi.org/10.17044/SCILIFELAB.25593405.

## Author contributions

N.S. and L.S. conceived and designed the study. N.S. and H.T. conducted experiments. N.S. analyzed data and drafted the manuscript, and all authors revised and edited the manuscript. L.S. secured funding and supervised the project.

## Supporting information

Figure S1. Nutrient levels during phenotype transition on 0.5xTSA.

Figure S2. Influence of nutrients and developmental mutations on foraging phenotypes.

Figure S3. Foraging by *Streptomyces* species.

Figure S4. Mold inhibition and metabolomics of foraging *S. coelicolor*.

Table S1. Prediction of signal peptides in the foraging secretome using SignalP 6.0.

Table S2. PANTHER classification of signal peptide containing proteins with regard to protein class.

Table S3. TTA containing genes in *S. coelicolor* A3(2).

Table S4. Mass spectrometric identification of locally isolates streptomycetes.

Table S5. Metabolomic features annotated using monoisotopic mass and StreptomeDB.

Table S6. Monoisotopic mass of unannotated features significantly higher levels in FAM conditions. Table S7. Strain list.

Video S1. *S. coelicolor* phenotype transition.

Video S2. *S. coelicolor* growing on FAM-agar.

Video S3. Addition of glucose to foraging *S. coelicolor*.

Video S4. Addition of tryptone to foraging *S. coelicolor*.

## Supporting figure legends

**Figure S1.** Nutrient levels during phenotype transition on 0.5xTSA. (**A**) The fold change for every nutrient at every time point is plotted for both sampled positions on the dish. The plot reveals a highly positive correlation, as indicated by the R-value of >0.96 suggesting a significant linear relationship. (**B**) The fold change for each annotated metabolite during phenotype transition on 0.5xTSA is overlayed for both positions on the dish. The time point where the transition was observed is indicated with the dotted lines at day 21.

**Figure S2.** Influence of nutrients and developmental mutations on foraging phenotypes. (**A**) Micrograph showing the phenotypic growth on WT *S. coelicolor* inoculated on FAM, FAM+sucrose (+S), FAM+glucose (+G) and FAM+fructose (+F) for 15 weeks at 30°C. (**B**) Growth of *S. coelicolor* wild-types (M145 and M600) and developmental mutants: *bldA*, *whiA* (J2401), *whiB* (J2402), *whiG* (J2400), *whiH* (J2408), *whiI* (J2450), and M851 (Δ*adpA*) on FAM.

**Figure S3.** Foraging by *Streptomyces* species. **(A)** Sector point mutations of foraging colonies. Spores of *S. coelicolor* were inoculated on FAM-agar and the sectors were isolated and subjected to Illumina sequencing. The inoculum was compared to the first level sector formed (α) and two succeeding sectors (I and IV) originating from lineage α. Using the breseq software, two point-mutations were detected in sector α: DnaA (SCO3879:E520K) and uncharacterized DUF349-containing protein (SCO1511:+G). One mutation was found in SCO5585 (CG)5 Ò 4 in linage IV. (**B**) Streptomycetes on FAM stained with Lugol’s solution after 5 days of incubation. Abbreviations: SANT; *S. antibioticus*, SANU; *S. anulatus*, SCO; *S. coelicolor*, SFI; *S. filipinensis*, SGR; *S. griseus*, SGRI; *S. griseolus*, SHIR; *S. hiroshimensis*, SHYG; *S. hygroscopicus*, SKAN; *S. kanamyceticus*, SLIV; *S. lividans*, SNOR; *S. norsei*, SVEN; *S. venezuelae*, KSET; *Kitasatospora setae*. (**C**) Micrographs of the tested streptomycetes grown on FAM+G for 14 days. (**D**) Micrographs of the tested streptomycetes grown on FAM+T for 14 days.

**Figure S4.** Mold inhibition and metabolomics of foraging *S. coelicolor*. (**A**) Inhibition of *P. citrinum by S. coelicolor* developmental mutants: *whiA* (J2401), *whiB* (J2402), *whiG* (J2400), *whiH* (J2408), *whiI* (J2450) and wild-type strain M600. (**B**) Inhibition of local *Aspergillus*-isolate in an overlay-assay on foraging *S. coelicolor*. Pictures were taken of foraging colonies using a dissection microscope. (**C**) Graphs showing the features annotated by monoisotopic mass and StreptomeDB. All graphs depict statistically significant outcomes comparing FAM and FAM+G as determined by t-tests, with p-values <=0.0002. 7-Acetyl-3.6-dihydroxy-8-methyl-tetralone is abbreviated as 7A36D8MT in the plot. (**D**) Venn-diagram showing the number of shared and unique metabolites detected during growth on FAM and FAM+G. **(E**) Plots of the unannotated metabolomic features unique to growth on FAM. The top right corner of each graph indicates the molecular weight of corresponding metabolite. All graphs depict statistically significant outcomes as determined by t-tests, with p-values <0.0001.

