## Supplementary figures and images for "Nutrient deprived growth of *Streptomyces* promotes foraging growth and enhanced antimicrobial activity"

### Supplementary figures

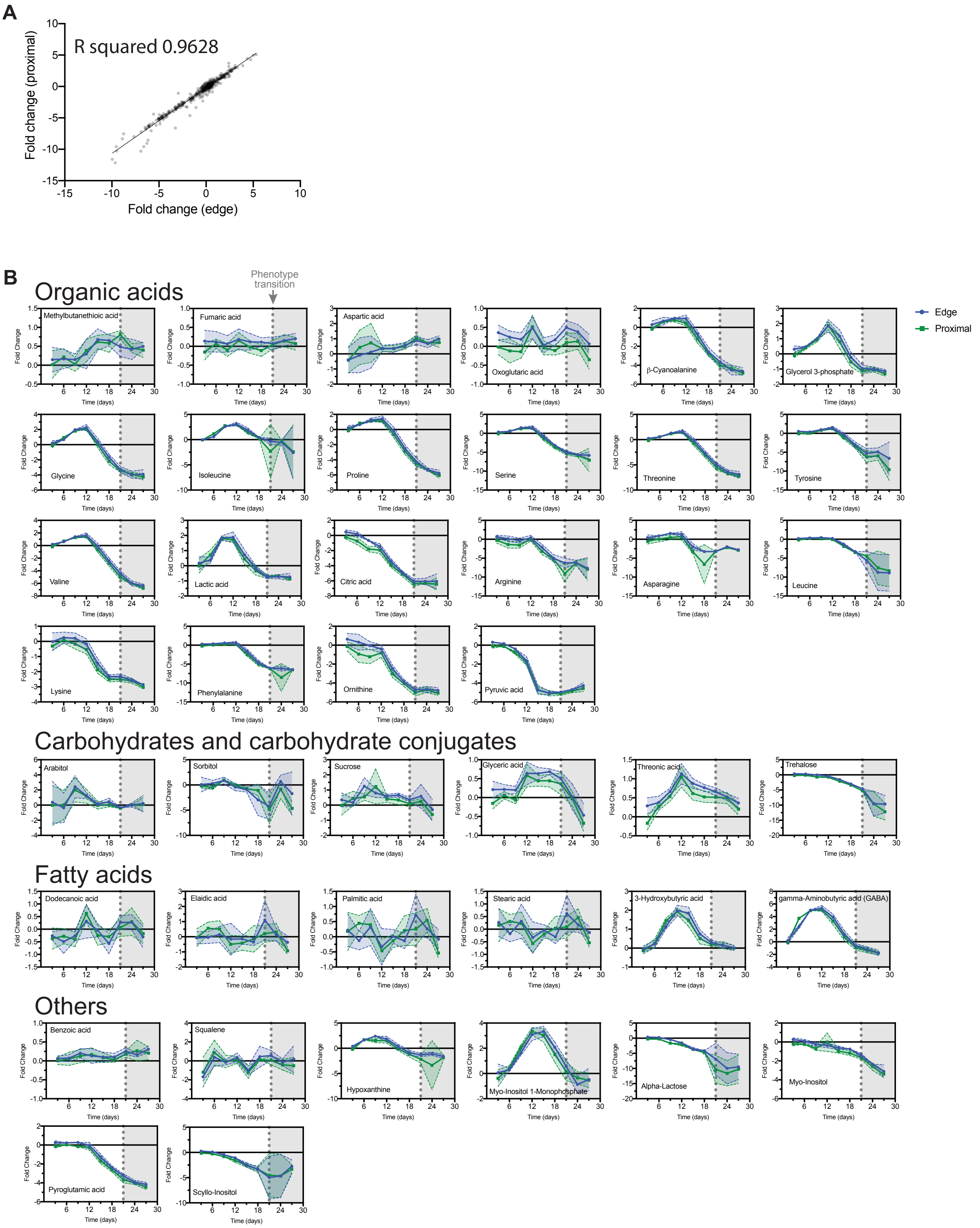

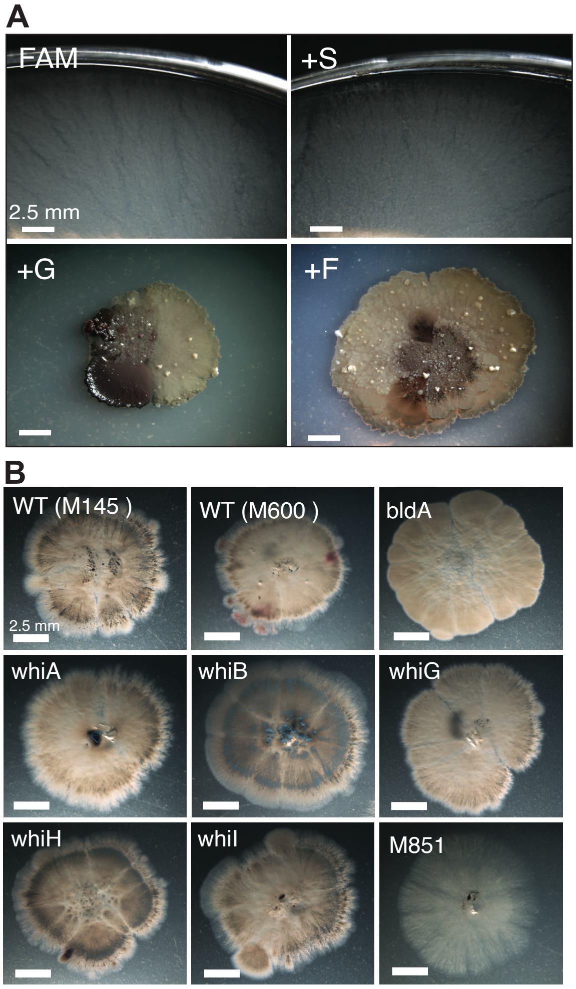

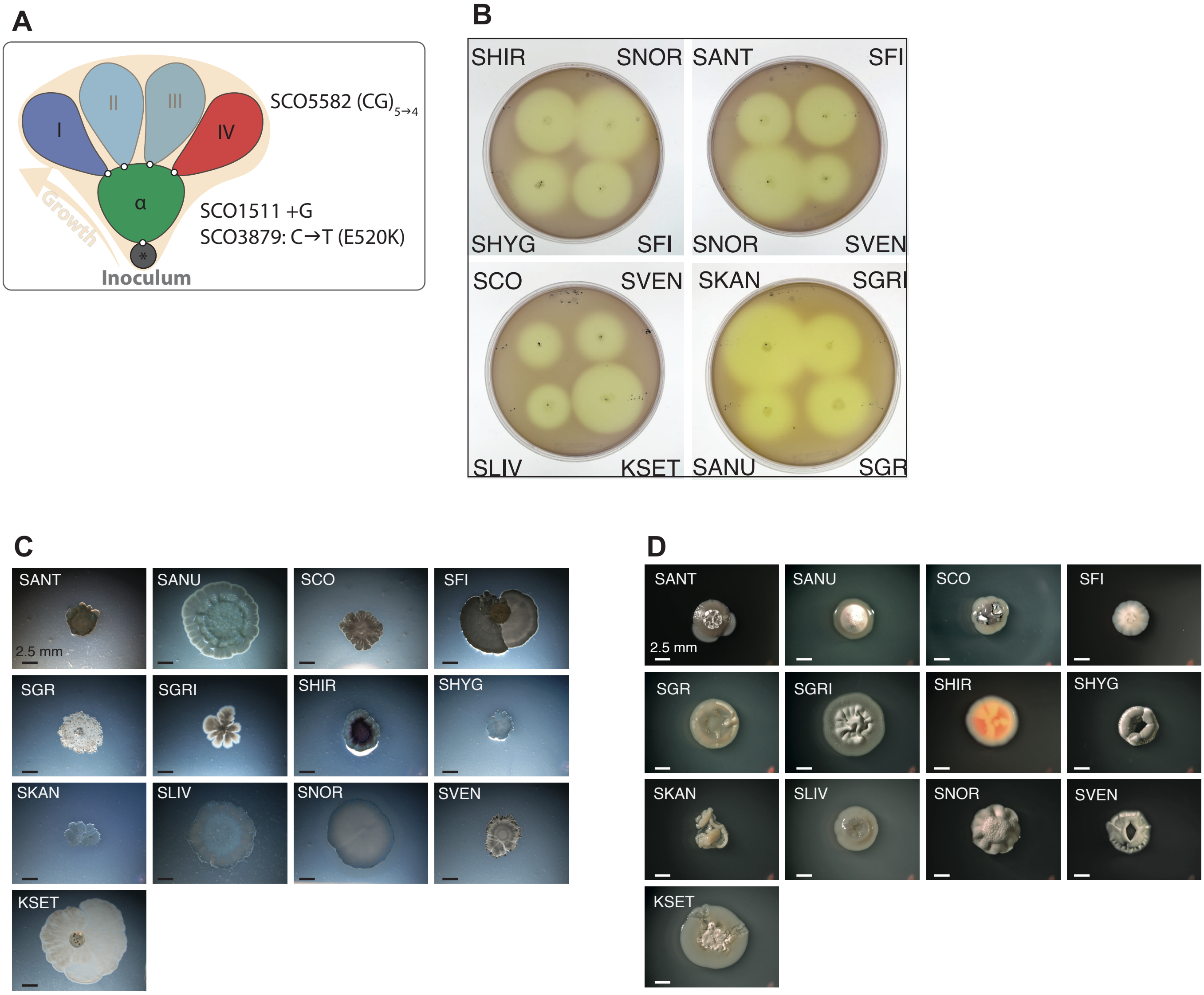

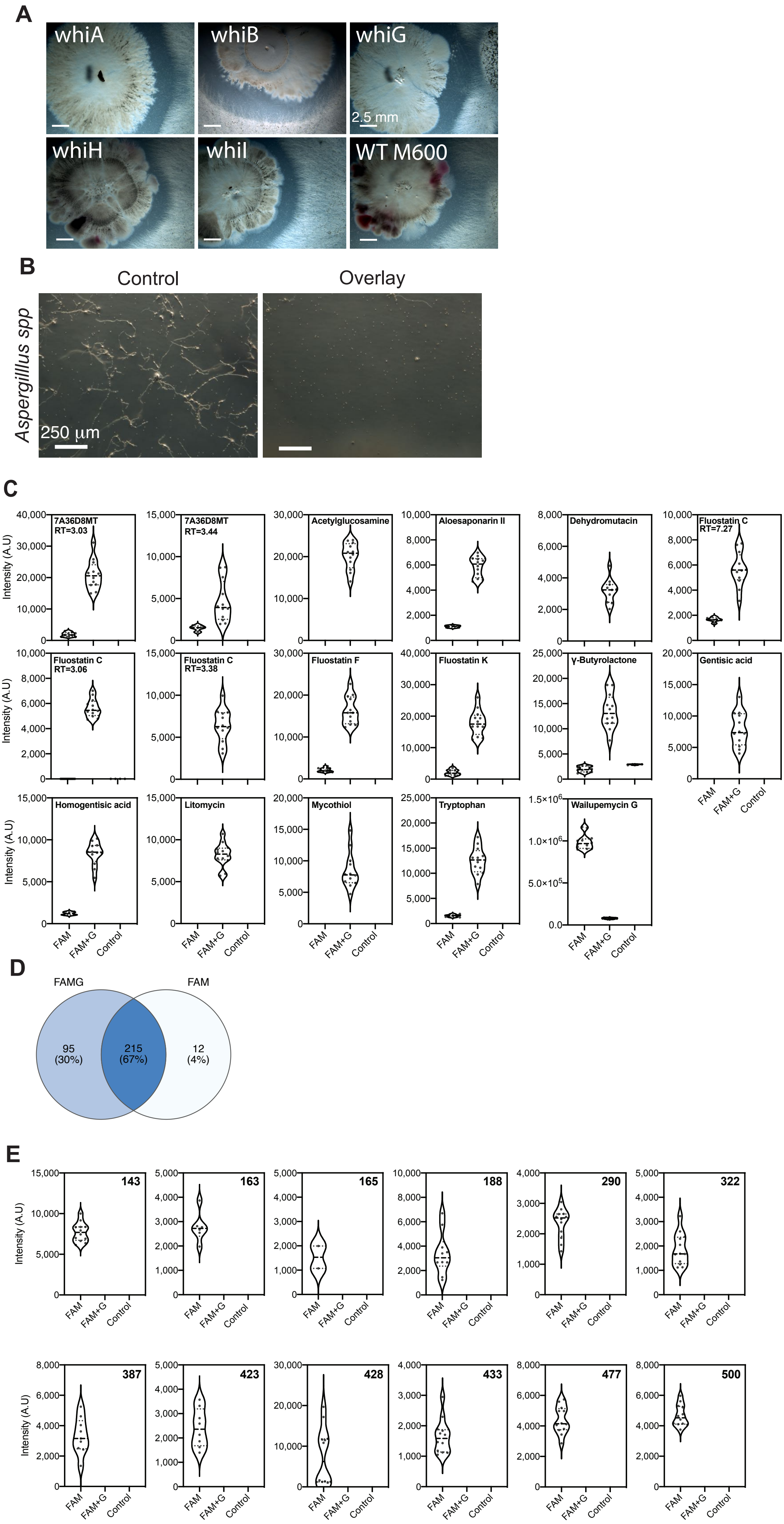
